## Supplemental Information for "A lncRNA identifies *IRF8* enhancer element in negative feedback control of dendritic cell differentiation"

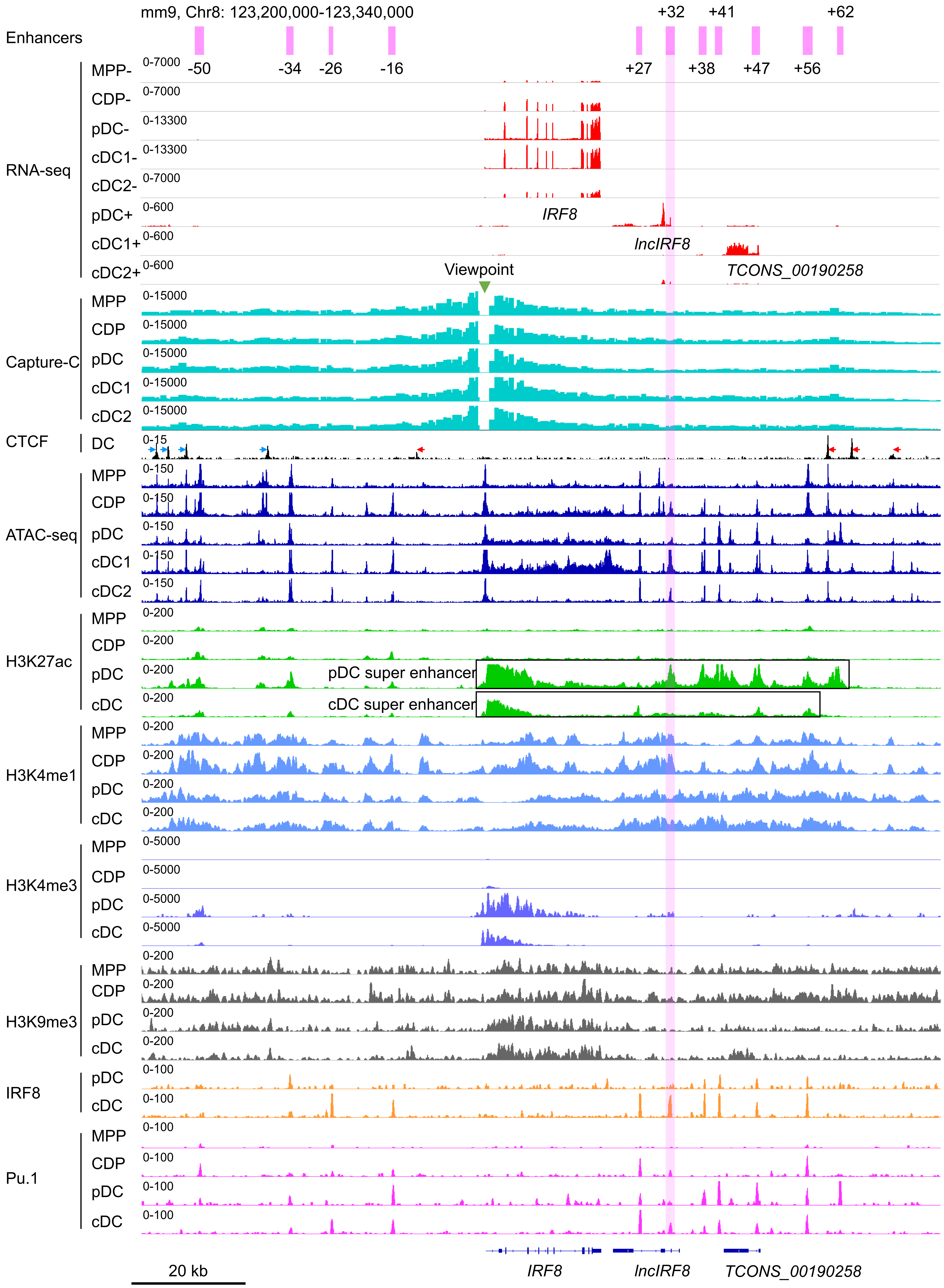


**Figure S1. Related to Figure 1. Epigenetic signatures of *IRF8* locus during DC differentiation**

Gene expression and epigenetic signatures of *IRF8* gene and its flanking regions in MPP, CDP, pDC, all cDC, cDC1 and cDC2 are visualized by IGV browser. Gene expression by RNA-seq, chromatin interactions with *IRF8* promoter by Capture-C, chromatin accessibility by ATAC-seq, histone modification (H3K27ac, H3K4me3, H3K4me1, H3K9me3) and TF binding (CTCF, IRF8 and PU.1) by ChIP-seq are as in Figure 1A and B. Positions of upstream and downstream *IRF8* enhancers are in purple, Capture-C viewpoint (green triangle), *IRF8* gene, pDC specific *lncIRF8* and cDC1 specific *TCONS_00190258* lncRNA are indicated. The orientation of CTCF binding is indicated with blue and red arrows. pDC and cDC super enhancers (Grajales-Reyes et al., 2015) are indicated with black boxes. For RNA-seq – and + strands are shown. Scale bar: 20 kb.

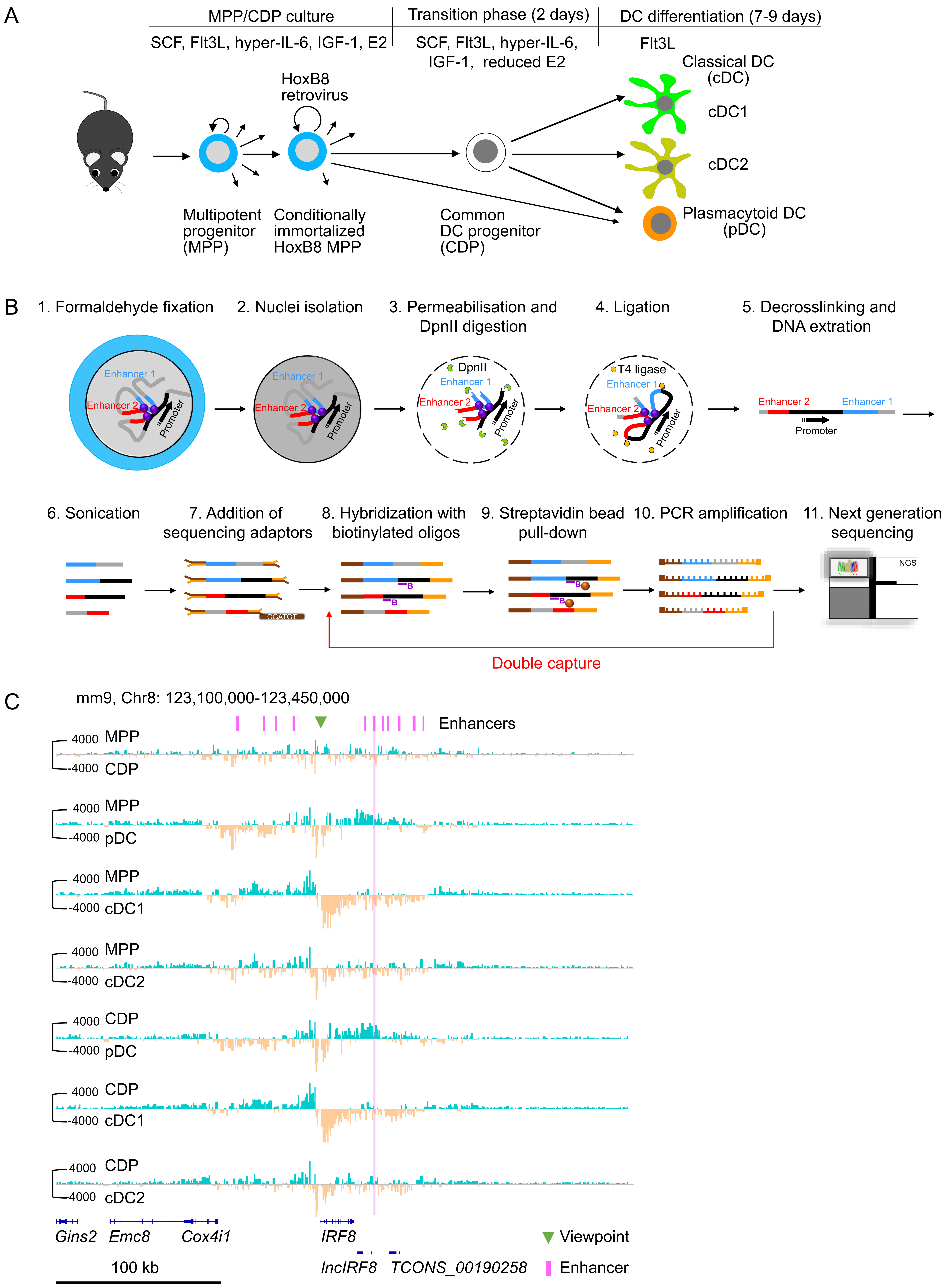


**Figure S2. Related to Figure 1. *In vitro* DC differentiation of HoxB8 MPP and nuclear-titrated (NuTi) Capture-C**

(A) HoxB8 MPP were obtained by infection of bone marrow MPP with HoxB8 retrovirus and cultured in growth medium with SCF, Flt3L, hyper-IL-6, IGF-1 and β-estradiol (E2) (MPP/CDP culture). DC differentiation is induced in growth medium plus high concentration of Flt3L and reduced E2 for 2 days (transition phase). Then growth factors and E2 are removed and cells are further differentiated into DC with high concentration of Flt3L only (DC differentiation).

(B) For nuclear-titrated (NuTi) Capture-C FACS sorted cells were fixed with formaldehyde and nuclei were isolated (step 1 and 2). Nuclei were permeabilized with SDS and digested with DpnII (step 3). DNA fragments in nuclei were re-ligated by T4 ligase (step 4) and DNA was extracted to obtain 3C libraries (step 5). The 3C libraries were subjected to sonication (~200 bp fragments), sequencing adaptors were added and DNA was then hybridized with 70 bp biotinylated oligos of the viewpoint, followed by streptavidin bead pull-down and PCR amplification (steps 6-10). Steps 8-10 were repeated to increase capture efficiency. Finally, the captured DNA was subjected to next-generation sequencing (step 11). This scheme was adapted from (Downes et al., 2022).

(C) Comparison of the chromatin interactions with the *IRF8* promoter by Capture-C (viewpoint, green triangle) between MPP, CDP, pDC, cDC1 and cDC2. Positions of *IRF8* enhancers are indicated (purple). Pairwise comparisons are shown and color coded (turquoise vs orange) as in Figure 1C. Differential tracks were created by subtraction of the mean normalized tracks of Figure 1B. Scale bar: 100 kb.

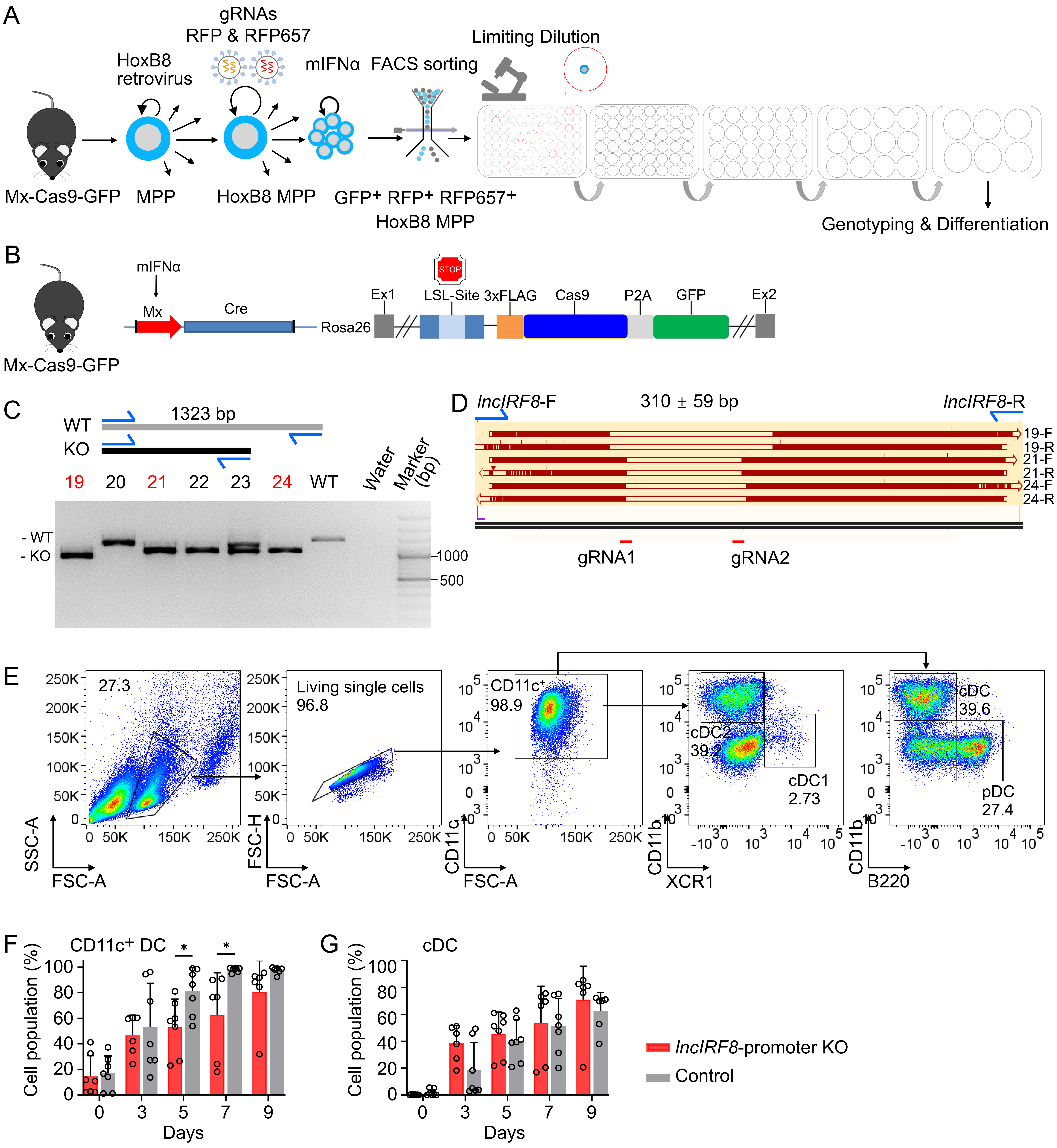


**Figure S3. Related to Figure 2. Generating *lncIRF8* promoter KO HoxB8 MPP and its impact on DC differentiation**

(A) Mx-Cas9-GFP HoxB8 MPP were infected with 2 gRNA in RFP and RFP657 lentiviral vectors. Mouse interferon α (mIFNα) was applied for 4 hours to induce Cas9 and GFP expression. Cells were cultured for another 2 days and FACS sorted for GFP+ RFR+ RFP657+ cells and 3,000 FACS sorted HoxB8 MPP were subjected to limiting dilution to obtain single cell colonies. Single cell HoxB8 MPP were genotyped by genomic PCR and clones with homozygous deletions in the *lncIRF8*-promoter were used for further studies.

(B) Mx-Cas9-GFP mice contain of a mIFNα inducible Cre recombinase that removes the stop cassette and allows Cas9 and GFP expression. Cas9 and GFP protein are separated by the 2A self-cleaving peptide (P2A) and thus all GFP+ cells express Cas9.

(C and D) Representative genotyping of *lncIRF8*-promoter KO in single-cell Mx-Cas9-GFP HoxB8 MPP. Homozygous deletions in *lncIRF8*-promoter (red) were detected by PCR and agarose gel electrophoresis (C). *lncIRF8* promoter deletions determined by Sanger sequencing in three representative KO clones of (C) are depicted in (D). *lncIRF8*-F and *lncIRF8*-R refers to primers for genotyping and Sanger sequencing, and sequences for both directions, referred to F and R, of clones 19, 21 and 24 are shown. Positions of gRNA1 and gRNA2, which were used for generating the deletion in the *lncIRF8*-promoter, are indicated.

(E) Gating strategy for Flt3L directed DC differentiation. CD11c+ DC were gated on CD11b- B220+ pDC, CD11b+ B220- cDC, CD11blow/- XCR1+ cDC1, and CD11b+ XCR1- cDC2.

(F-G) Quantification of CD11c+ DC and all cDC of panel (E) of living single cells in *lncIRF8*-promoter KO HoxB8 MPP and control on Flt3L directed DC differentiation at day 0, 3, 5, 7, and 9. n=6-7. Data represent mean ± SD of at least 3 independent experiments with different HoxB8 MPP clones of *lncIRF8*-promoter KO and control. *$P$<0.05, multiple t-tests. Data that have no difference ($P$>0.05) are not labeled.

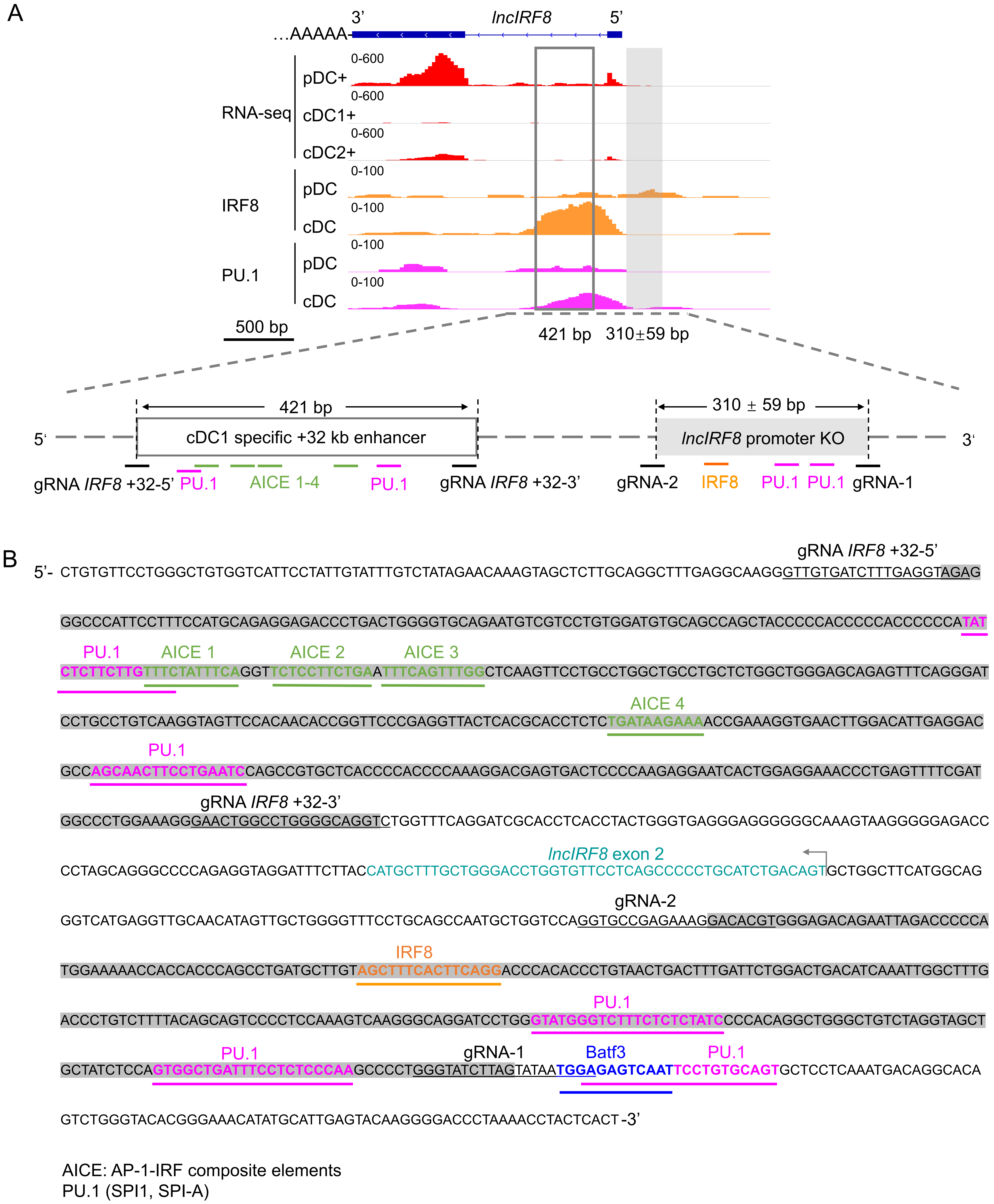


**Figure S4. Related to Figure 2. Information of *lncIRF8* promoter KO**

(A) Schematic representation of *lncIRF8* locus. The sequences deleted in the *lncIRF8* promoter KO are indicated by a grey filled box and multiple potential IRF8 and PU.1 binding sites are depicted. The cDC1 specific +32 kb enhancer by (Durai et al., 2019) is shown by an open box. Gene expression was by RNA-seq and IRF8 and PU.1 TF binding was by ChIP-seq as in Figure 1A, Figure 2A and Figure S1. Scale bar: 500 bp.

(B) Sequence of *lncIRF8* promoter and cDC1 specific +32 kb enhancer. Sequences underlined in black indicate gRNAs used for generating the deletions. Sequence motives for AP1-IRF composite elements (AICE), IRF8, PU.1, and Batf3 were indicated with green, orange, purple, and blue lines, respectively. *lncIRF8* exon 2 is in turquoise; arrow indicates *lncIRF8* transcriptional start site and direction. The shaded sequences represent the KO region of *lncIRF8* promoter and the cDC1 specific +32 kb enhancer by (Durai et al., 2019).

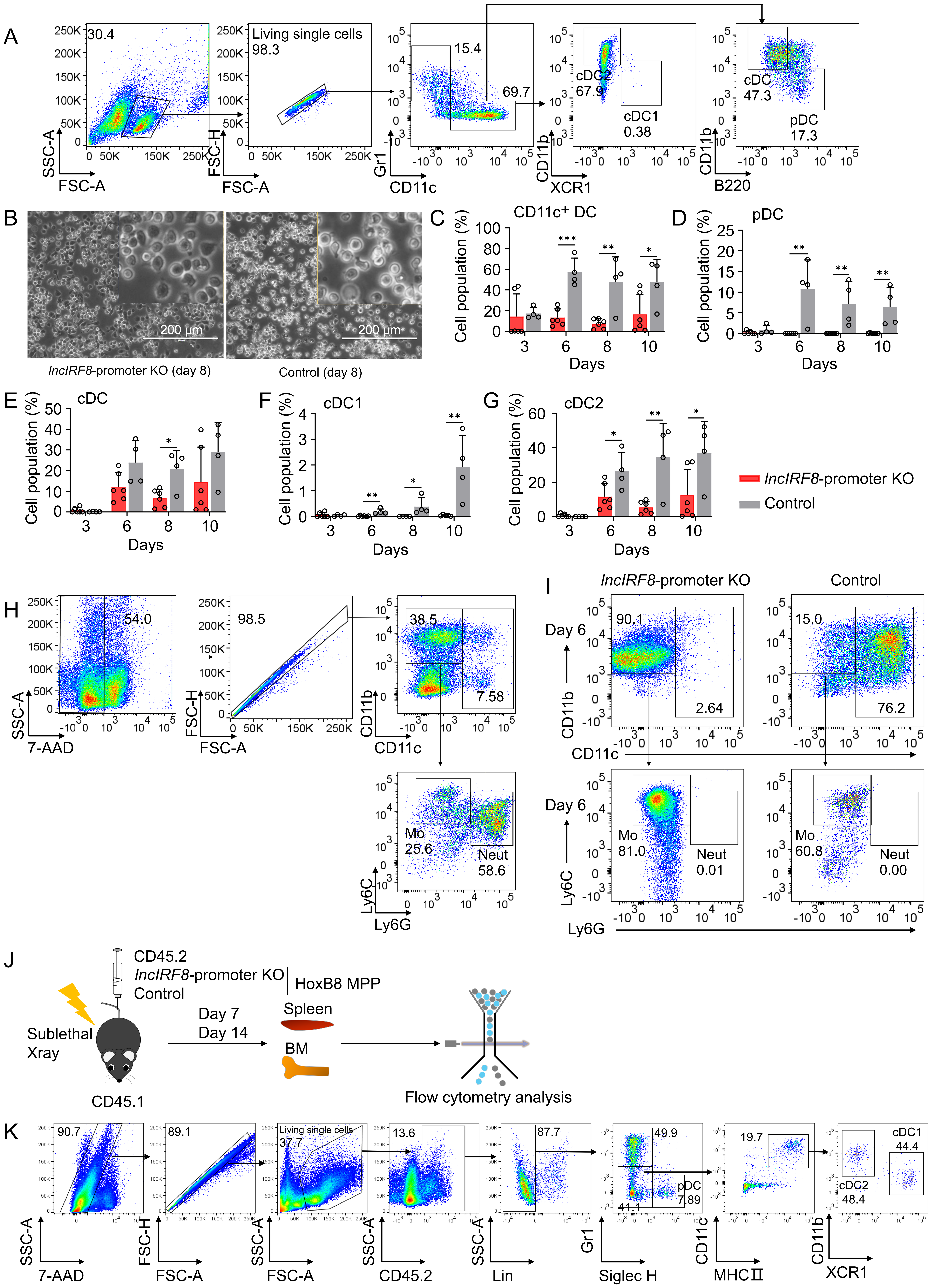

**Figure S5. Related to Figures 2 and 3. Impact of *lncIRF8* promoter KO on spontaneous DC differentiation and cell transplantation**

(A) Gating strategy for spontaneous DC differentiation. Gr1+ monocytes, and the pDC, all cDC, cDC1 and cDC2 subsets from Gr1- CD11c+ cells were gated as in Figure S3E.

(B) Representative phase-contrast microscopy images of *lncIRF8*-promoter KO HoxB8 MPP and control on spontaneous DC differentiation (-E2) day 8. Scale bar: 200 µm.

(C-G) Quantification of $CD11c^+$ DC, pDC, all cDC, cDC1 and cDC2 of panel (A) of living single cells in *lncIRF8*-promoter KO HoxB8 MPP and control on spontaneous DC differentiation (-E2) day 3, 6, 8, and 10. n=6 for *lncIRF8*-promoter KO; n=4 for KO control. Data represent mean ± SD of at least 3 independent experiments with different HoxB8 MPP clones of *lncIRF8*-promoter KO and control without deletion. $^*P<0.05$, $^{**}P<0.01$, $^{***}P<0.001$, multiple t-tests. Data that have no difference ($P>0.05$) are not labeled.

(H) Gating strategy for identification of $Gr1^+$ monocytes upon spontaneous DC differentiation in Figure 2E and Figure S5A. Mouse BM cells were used and $Gr1^+$ monocytes (Mo) were gated as 7-$AAD^-$ $CD11b^+$ $CD11c^-$ $Ly6C^+$ $Ly6G^-$. Neutrophils (Neut) were gated as 7-$AAD^-$ $CD11b^+$ $CD11c^-$ $Ly6C^{low/-}$ $Ly6G^+$.

(I) Representative flow cytometry analysis of $Gr1^+$ monocytes at day 6 of spontaneous DC differentiation of *lncIRF8*-promoter KO and control. Cells were gated as in panel (H).

(J) Workflow of HoxB8 MPP transplantation. CD45.1 recipient mice were sublethal irradiated and injected with CD45.2 *lncIRF8*-promoter KO and control HoxB8 MPP. Seven and 14 days after cell transplantation, mice were sacrificed and cells from bone marrow and spleen were isolated and subjected to flow cytometry analysis for DC subsets.

(K) Gating strategy for CD45.2 DC subsets following transplantation of CD45.2 *lncIRF8*-promoter KO HoxB8 MPP into CD45.1 recipient mice. 7-$AAD^-$ $CD45.2^+$ $lineage^-$ donor cells were gated on $Gr1^+$ monocytes, $Gr1^-$ Siglec $H^+$ pDC, and $Gr1^-$ Siglec $H^-$ $CD11c^+$ $MHCII^+$ DC, and further on cDC1 and cDC2 subsets. cDC1 are $Gr1^-$ Siglec $H^-$ $MHCII^+$ $CD11c^+$ $CD11b^{low/-}$ $XCR1^+$ and cDC2 are $Gr1^-$ Siglec $H^-$ $MHCII^+$ $CD11c^+$ $CD11b^+$ $XCR1^-$. The lineage cocktail (Lin) was: Ter119, CD19, CD3e, NK1.1, F4/80.

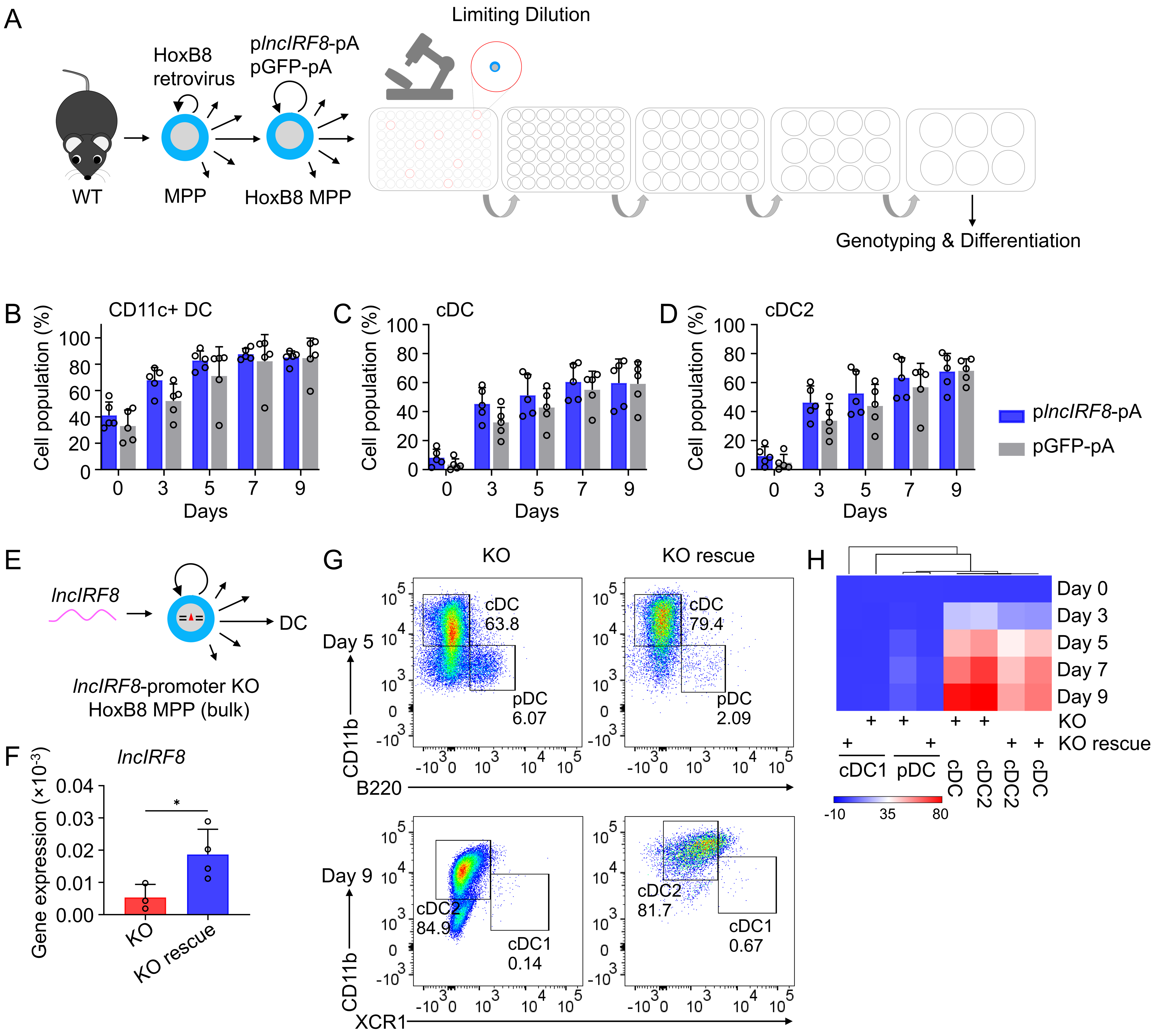


**Figure S6. Related to Figure 4. *lncIRF8* overexpression and rescue left DC differentiation unaffected**

(A) WT HoxB8 MPP were infected with p*lncIRF8*-pA (*lncIRF8* overexpression) and pGFP-pA (control) lentiviral particles, followed by limiting dilution to obtain single cell colonies. HoxB8 MPP clones were screened by genomic PCR and the positive colonies were subjected to DC differentiation.

(B-D) Quantification of CD11c+ DC, all cDC and cDC2 of living single cells in p*lncIRF8*-pA and pGFP-pA HoxB8 MPP on day 0, 3, 5, 7, and 9 of Flt3L directed DC differentiation (n=5). Gating was as in Figure S3E. Data represent mean ± SD of at least 3 independent experiments with different HoxB8 MPP clones of p*lncIRF8*-pA and pGFP-pA. Data were calculated by multiple t-tests and data that have no difference ($P$>0.05) are not labeled.

(E) Schematic representation of *lncIRF8* rescue in *lncIRF8*-promoter KO HoxB8 MPP (bulk).

(F) Gene expression of *lncIRF8* in *lncIRF8*-promoter KO and KO rescue HoxB8 MPP. Gene expression was by RT-qPCR and normalized to *GAPDH.* Data represent mean ± SD of at least 3 independent experiments and were calculated by unpaired t test. *$P$<0.05.

(G) Representative flow cytometry analysis of DC subsets pDC, cDC1 and cDC2 at day 5 and 9 of Flt3L directed DC differentiation of *lncIRF8*-promoter KO rescue and control (KO).

(H) DC subsets pDC, all cDC, cDC1 and cDC2 were calculated in percent of living single cells in *lncIRF8*-promoter KO and KO rescue on day 0, 3, 5, 7, and 9 of Flt3L directed DC differentiation and data are represented in heat map format; n=3-4 from at least 3 independent experiments. Red, high frequency; white, intermediate frequency and blue, low frequency.

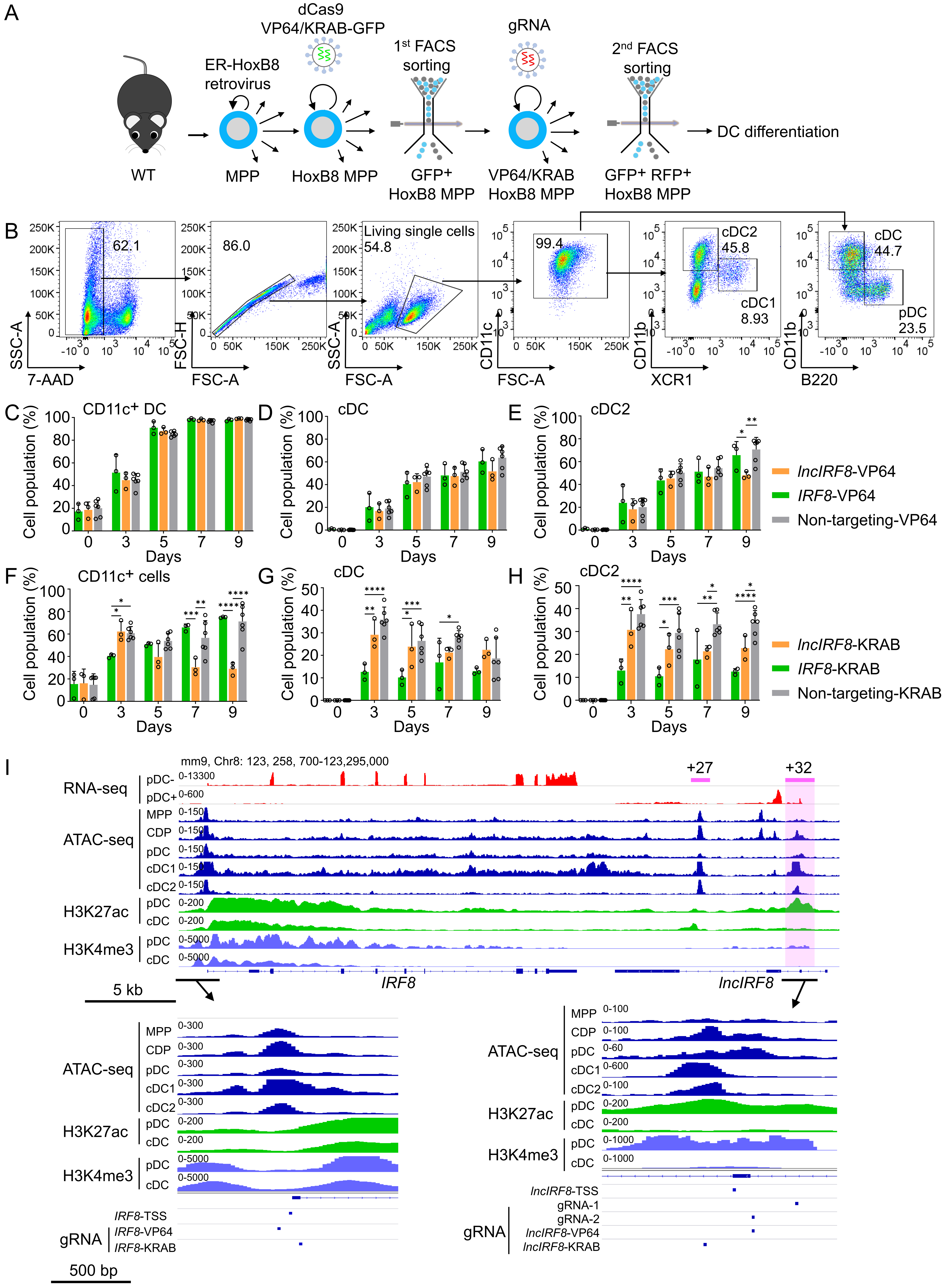


**Figure S7. Related to Figures 2, 5 and 6. CRISPR/dCas9 mediated gene activation and repression, and gRNA positions**

(A) WT HoxB8 MPP were infected with dCas9-VP64-GFP and KRAB-dCas9-GFP lentiviral vectors followed by 1st FACS sorting for GFP+ cells to obtain dCas9-VP64-GFP and dCas9-KRAB-GFP HoxB8 MPP. Gene targeting of dCas9-VP64 and dCas9-KRAB was achieved by infection of MPP with specific gRNA vectors and 2nd FACS sorting for GFP+ RFP+ cells.

(B) Gating strategy for DC subsets in CRISPR/dCas9 HoxB8 MPP upon DC differentiation. DC subsets were gated from 7-AAD- CD11c+ cells, and further gated on all cDC, cDC1, cDC2 and pDC as in Figure S3E.

(C-H) Quantification of $CD11c^{+}$ cells, all cDC, and cDC2 of panel (B) in percent of living single cells in dCas9-VP64 cells and dCas9-KRAB cells on day 0, 3, 5, 7, and 9 of Flt3L directed DC differentiation (n=3). Non-targeting-VP64 and Non-targeting-KRAB refers to non-targeting-VP64 and KRAB controls, respectively (n=6).

Data represent mean ± SD of 3 independent experiments. *$P<0.05$, **$P<0.01$, ***$P<0.001$, ****$P<0.0001$, two-way ANOVA, Tukey's multiple comparisons test. Data that have no difference ($P>0.05$) are not labeled.

(I) Gene expression and epigenetic landscape of *IRF8* gene and its flanking regions in MPP, CDP, pDC, all cDC, cDC1 and cDC2 are depicted as in Figure 1A and Figure S1. *IRF8* +27 kb and +32 kb enhancer (purple line and box, respectively), transcription start sites of *IRF8* (*IRF8*-TSS) and *lncIRF8* (*lncIRF8*-TSS) are indicated. Bottom, positions of gRNAs used in this study: gRNA1 and gRNA2, *lncIRF8* promoter KO by the CRISPR/Cas9 system; *IRF8*-VP64 and KRAB gRNAs for activation and repression of *IRF8* promoter, respectively; *lncIRF8*-VP64 and KRAB gRNAs for activation and repression of *lncIRF8* promoter, respectively.

**Table S1. Off-target analysis**

| Potential off-targets | | | KO bulk (100 cells) | KO single-cell clones | | | | | Gene expression | | | |
|---|---|---|---|---|---|---|---|---|---|---|---|---|
| | | | | 6 | 7 | 19 | 21 | 24 | MPP | CDP | cDC | pDC |
| gRNA1 | Chr 13: -48032179 | *GM36101* | No | No | No | No | No | No | No | No | No | No |
| | Chr 2: -131293076 | Non-coding | No | No | No | No | No | No | No | No | No | No |
| gRNA2 | Chr 1: -136050990 | *ASCL5* | 16 bp deletion | 16 bp deletion | 16 bp deletion | No | 1 bp insertion | No | No | No | No | No |
| | Chr 6: +119352090 | *CACNA2D4* | No | No | No | No | No | No | No | No | No | Yes |
| | Chr 3: +33982811 | Non-coding | 3-29 bp deletion; 1-3 bp insertion | 3-29 bp deletion | 3 bp deletion | 1-2 bp insertion | No | No | No | No | No | No |
| | Chr 2: -132060887 | Non-coding | No | No | No | No | No | No | No | No | No | No |
| | Chr 2: -150624255 | Non-coding | No | No | No | No | No | No | No | No | No | No |

Top potential off-targets of gRNA1 and gRNA2 predicted by CRISPR-Cas9 gRNA checker (see Methods Details) were analyzed in *lncIRF8* promoter KO bulk culture and KO single-cell clones by PCR analysis of genomic DNA. Potential off-target genes (coding) and non-coding sequences are listed. The absence of off-targets (No) and off-target deletions/insertion are shown.
