## Supplementary figures and images for "A lncRNA identifies *IRF8* enhancer element in negative feedback control of dendritic cell differentiation"

### Graphical Abstract

Xu et al., Graphical Abstract

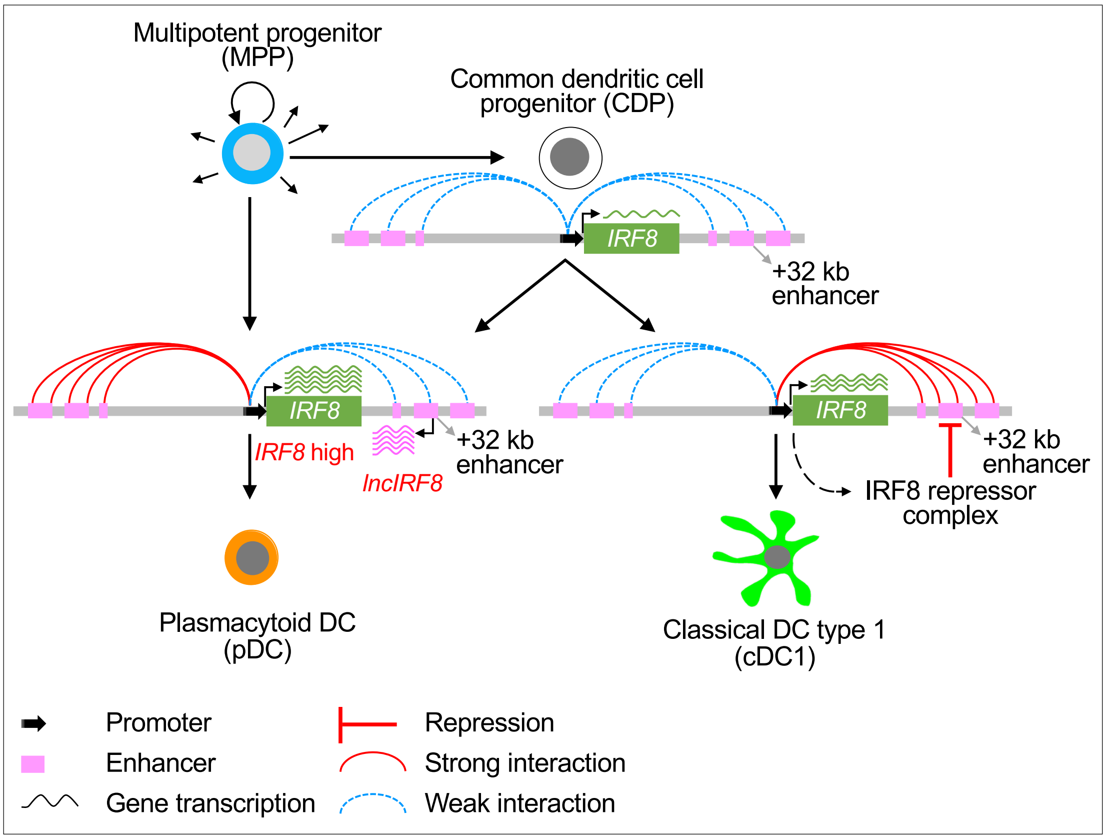
